# Targeting CHEK2-YBX1&YBX3 regulatory hub to potentiate immune checkpoint blockade response in gliomas

**DOI:** 10.1101/2025.03.09.642289

**Authors:** Heba Ali, Ningjia Zhou, Li Chen, Levi van Hijfte, Vivekanudeep Karri, Yalu Zhou, Karl Habashy, Victor A. Arrieta, Kwang-Soo Kim, Joseph Duffy, Ragini Yeeravalli, Deanna M. Tiek, Xiao Song, Snehasis Mishra, Catalina Lee-chang, Atique U. Ahmed, Dieter Henrik Heiland, Adam M. Sonabend, Crismita Dmello

## Abstract

Although GBM’s immunosuppressive environment is well known, the tumor’s resistance to CD8+ T cell killing is not fully understood. Our previous study identified Checkpoint Kinase 2 (Chek2) as the key driver of CD8+ T cell resistance in mouse glioma through an in vivo CRISPR screen and demonstrated that Chk2 inhibition, combined with PD-1/PD-L1 blockade, significantly enhanced CD8+ T cell-mediated tumor killing and improved survival in preclinical model. Here, we aimed to elucidate the immunosuppressive function of Chek2. Immunoprecipitation (IP) followed by mass spectrometry (MS) and phosphoproteomics identified an association between Chek2 with the DNA/RNA-binding proteins YBX1 and YBX3 that are implicated in transcriptional repression of pro-inflammatory genes. Single-gene knock-out and overexpression studies of CHEK2, YBX1, and YBX3 in multiple glioma cell lines revealed that these proteins positively regulate each other’s expression. RNA sequencing coupled with chromatin immunoprecipitation-sequencing (ChIP-seq) analysis demonstrated common inflammatory genes repressed by CHK2-YBX1&YBX3 hub. Targeting one of the hub proteins, YBX1, with the YBX1 inhibitor SU056 led to degradation of CHK2-YBX1&YBX3 hub. Targeting of this hub by SU056 led to enhanced antigen presentation and antigen specific CD8+ T cell proliferation. Further, combination of SU056 with ICB significantly improved survival in multiple glioma models. Collectively, these findings reveal an immunosuppressive mechanism mediated by the CHK2-YBX1&YBX3 hub proteins. Therefore, CHK2-YBX1&YBX3 hub targeting in combination with immune checkpoint blockade therapies in gliomas is warranted.

## Introduction

Glioblastoma (GBM) is an aggressive primary brain tumor characterized by rapid progression, therapy resistance, and a highly immunosuppressive tumor microenvironment (TME) (1,2). Despite multimodal treatment strategies, including surgical resection, radiotherapy, and temozolomide chemotherapy, patient prognosis remains poor, with a median survival of approximately 15 months (2,3). A major limitation in improving GBM therapy is its ability to evade immune detection and suppress anti-tumor immune responses (4–7). The GBM immune microenvironment is largely dominated by tumor-associated macrophages (TAMs) and myeloid-derived suppressor cells (MDSCs), which actively suppress cytotoxic T cell function, promote immunosuppression, and contribute to resistance against immune checkpoint blockade (ICB) therapy (8,9). Despite groundbreaking success of immune checkpoint inhibitors (ICIs) in cancers with high mutational burden and robust T-cell infiltration, their efficacy in gliomas remains limited (10). GBM employs various mechanisms to evade anti-tumor immunity, causing T-cell anergy, exhaustion, apoptosis, and sequestration in the bone marrow, posing significant challenges to immune-based approaches (11–15). Unlike other tumors with high mutation burden, such as melanoma or lung cancer, GBM is classified as an immunologically “cold” tumor, exhibiting limited T cell infiltration and a high degree of T cell dysfunction, making ICB therapies largely ineffective (4,16).

Our recent CRISPR-based kinome screen identified checkpoint kinase 2 (Human Checkpoint Kinase 2 gene/protein-*CHEK2*/CHK2; Mouse Checkpoint kinase 2 gene/protein*-Chek2*/Chk2) as a crucial regulator of immune suppression in glioma (17). CHEK2, a well-characterized DNA damage response (DDR) kinase, plays a key role in mediating cell cycle arrest and apoptosis in response to DNA damage (18,19). While traditionally recognized as a tumor suppressor, emerging studies indicate that *CHEK2* may also modulate innate immune signaling and inflammatory responses (20–24). Loss of *CHEK2* in glioma cells has been shown to enhance CD8+ T cell-mediated tumor killing, suggesting a role in immune evasion and checkpoint blockade resistance (17). Moreover, pharmacological inhibition of *CHEK2* augments type I interferon signaling and antigen presentation, reinforcing its relevance in tumor-immune interactions.

Beyond classical DDR components, RNA-binding proteins (RBPs) are increasingly recognized as hubs in immune regulation (25,26). Among them, Y-box binding protein 1 (YBX1) and YBX3 play pivotal roles in post-transcriptional gene regulation, inflammatory signaling, and stress response adaptation (27,28). YBX1 is overexpressed in multiple cancers, including GBM, where it modulates gene expression programs that drive tumor progression and therapy resistance (27,29). It has been implicated in regulating immune signaling pathways, including interferon responses and cytokine expression, though its precise role in GBM immunosuppression remains incompletely understood. Similarly, YBX3, another member of the Y-box protein family, has been linked to immune homeostasis, inflammation, and tumor microenvironment remodeling, but its functions in GBM remain largely unexplored (30). Given their ability to regulate immune-related gene expression networks, targeting YBX proteins may provide novel therapeutic opportunities in GBM.

In this study, we extend our previous findings by characterizing the molecular mechanisms through which Chk2 protein mediates immune suppression. Immunoprecipitation followed by mass spectrometry and phosphoproteomic analyses revealed that Chk2 interacts with RNA-binding proteins YBX1 and YBX3, key regulators of immune-related gene expression. Pharmacological inhibition using the YBX1 inhibitor SU056 effectively suppressed CHK2, YBX1 and YBX3 protein levels, leading to the restoration of pro-inflammatory pathways like antigen presentation and response to PD-1 and PD-L1 ICI therapy. This study establishes a mechanistic framework for protein hub driven immune modulation and offers a promising therapeutic strategy to potentiate ICI therapy for highly immunosuppressive tumors like GBM.

## Results

### Glioma-intrinsic Chk2 interacts with DNA- and RNA-binding proteins

To investigate the immunomodulatory function of glioma-intrinsic CHK2, we overexpressed mouse *Chek2*-Flag in GL261 glioma cells (Fig. 1A). GL261 *Chek2*-Flag-overexpressing cells were orthotopically implanted into C57BL/6 mice, and tumors were harvested at the experimental endpoint (Fig. 1B). Tumor proteins were extracted from the brain tumor tissue, followed by immunoprecipitation (IP) using Flag antibody (Ab) and IgG Ab as a control. Mass spectrometry (MS) analysis of the IP fraction identified Chk2 as the second most abundant protein (log₂ fold change >1,-adjusted log_10_(p-value) *> 1.0), confirming the specificity of the IP experiment. Among the top interactors, vesicular glutamate transporter 2 (*Slc17a6), a key player in glutamate transport, and DmX-like protein 2 (*DMXL2*), a synaptic vesicle scaffold protein, were identified. Notably, DNA- and RNA-binding proteins YBX3 (log₂ fold change=1.86,-adjusted log_10_(p-value)*=*1.13) and YBX1 (log₂ fold change=1.54,-adjusted log10(p-value)=0.29) although not significant were found to interact with Chk2 (Fig. 1C). The presence of these Y-box proteins in the Chk2 interactome suggests a role of Chk2 in glioma cells through RNA and DNA-binding protein interactions. To assess the relevance of these interactions in human GBM, we compared The Cancer Genome Atlas (TCGA) GBM RNA-seq data with MS hits (log₂ fold change>1.5) from IP-MS analysis (Fig. 1D). A significant positive correlation was observed between *CHK2* and *YBX1* expression in TCGA GBM datasets (Spearman *r* = 0.50, *P* = 1.07 × 10⁻¹⁰; Fig. 1E), further supporting an association. To characterize CHK2-dependent phosphorylation substrates, we performed global phosphoproteomics in mouse GL261 glioma cells with non-targeting control (NTC) and *Chek2* knockout (KO) cells (Fig. 1F). Phospho-enrichment analysis identified FK506-binding protein 15 (FKBP15), a regulator of cytoskeletal organization, and Acinus (*Acin1*), a nuclear protein involved in RNA processing and apoptosis, as the topmost phosphorylated proteins in *Chek2* KO cells. Notably, phosphorylation at the S52 residue of YBX3 was significantly increased in *Chek2* KO cells compared to NTC cells (Fig. 1G), supporting the MS findings in (Fig. 1C). Co-immunoprecipitation (co-IP) assays (Fig. 1H) further validated a direct interaction between CHK2 and YBX3, whereas no detectable interaction was observed between CHK2 and YBX1 (Fig. 1I).

**Figure 1.**
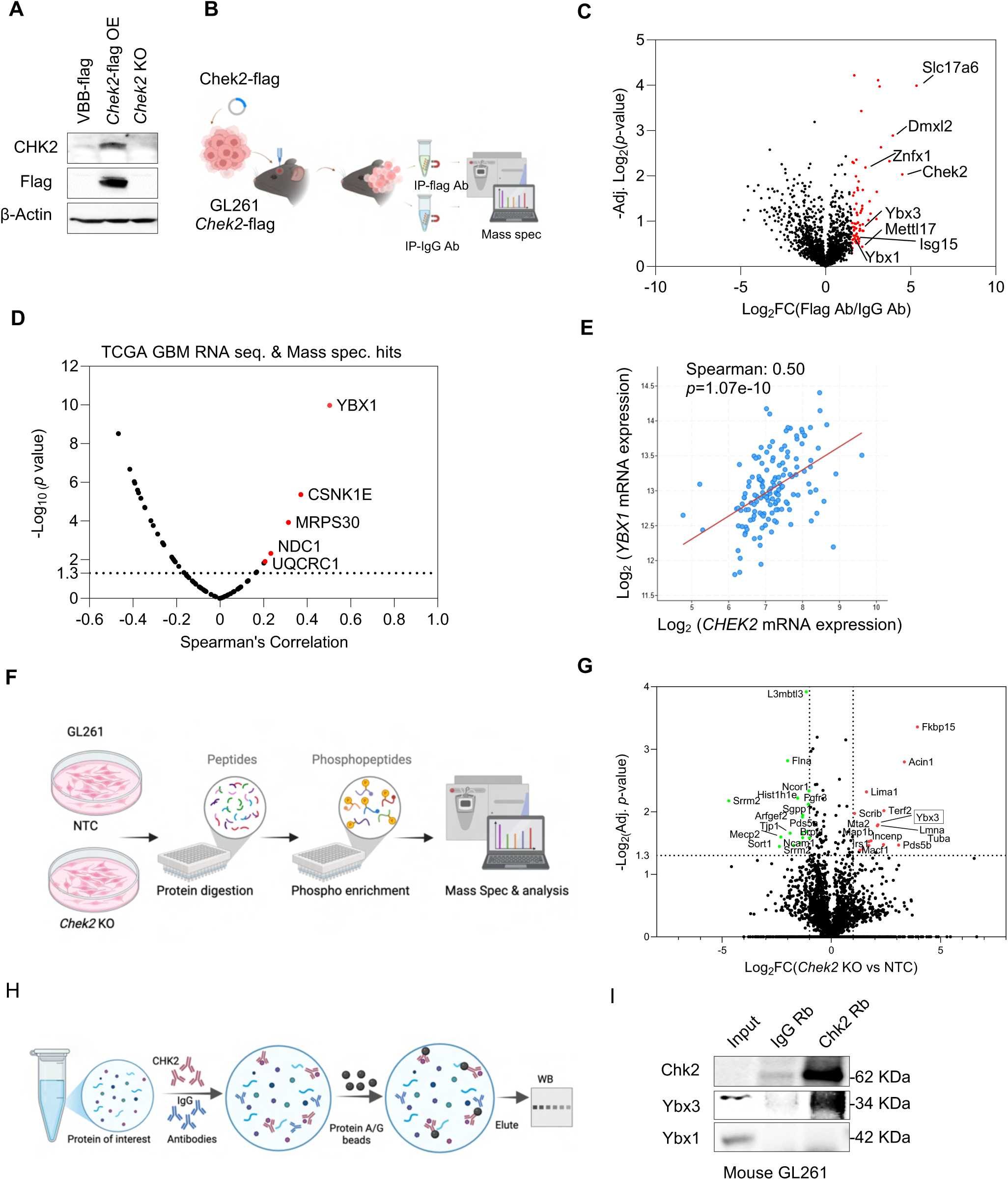
Glioma-intrinsic Chk2 interacts with DNA- and RNA-binding proteins. (A) Immunoblots of overexpression of FLAG-tagged Chek2 in mouse glioma cells GL261. (B) Glioma cells GL261 overexpressing Chek2 were injected intracranially into mice. Tumors were collected for immunoprecipitation followed by mass spectrometry analysis. (C) Volcano plot of mass spectrometry analysis of protein enrichment by immunoprecipitation. The plot displays the log2 fold-change (x-axis) versus the significance (-log10 p-value) for all proteins. Proteins shown in red represent the significantly increased proteins (log2 fold-change.1.5). (D) Volcano plot of TCGA GBM6 RNA-seq data integrated with mass spectrometry-based protein enrichment analysis. (E) Scatter plot showing the correlation between YBX1 and CHEK2 mRNA expression in the TCGA GBM dataset. Statistical analysis was performed using a two-tailed Pearson correlation, revealing a significant positive correlation (r = 0.50, P = 1.07 × 10⁻¹⁰). (F) Schematic representation of the phosphoproteomic assay. (G) Volcano plot showing log2 fold-change (average *Chek2* KO/average NTC) versus −log10 (p-value) for all quantified phosphopeptides. Proteins above the dashed line (> 1.3) on the y-axis are statistically significant. Red and green markers represent significantly increased and decreased in phosophorylation, respectively. (H-I) Schematic representative of co-immunoprecipitation assay followed by immunoblots of Chk2, Ybx1 and Ybx3 in mouse glioma cell line GL261.

### Positive feedback regulation between *CHK2*, *YBX1 and YBX3* in gliomas

We used gene knockout (KO) approaches in multiple GBM cell lines to dissect the interplay among *CHEK2*, YBX1, and YBX3. In U87 human GBM cells, *CHEK2* KO downregulated *YBX3* protein levels and did not significantly alter YBX1 levels (Fig. 2A). However, *YBX1* KO led to a marked reduction in *CHEK2* and YBX3 protein levels, and *YBX3* KO downregulated *CHEK2 levels but upregulated YBX1 levels.* In GBM6 human GBM cells *CHEK2* KO resulted in a significant reduction in both *YBX1* and *YBX3* protein levels (Fig. 2B). GBM6 *YBX1* KO reduced *CHEK2* and *YBX3 levels*. *YBX3* KO led to decreased *CHEK2* expression but triggered a compensatory upregulation of *YBX1*. Of note the guides sequences used for generation of CRISPR KO were unique to that specific gene and had no sequence similarity to other two genes that it regulated.

**Figure 2:**
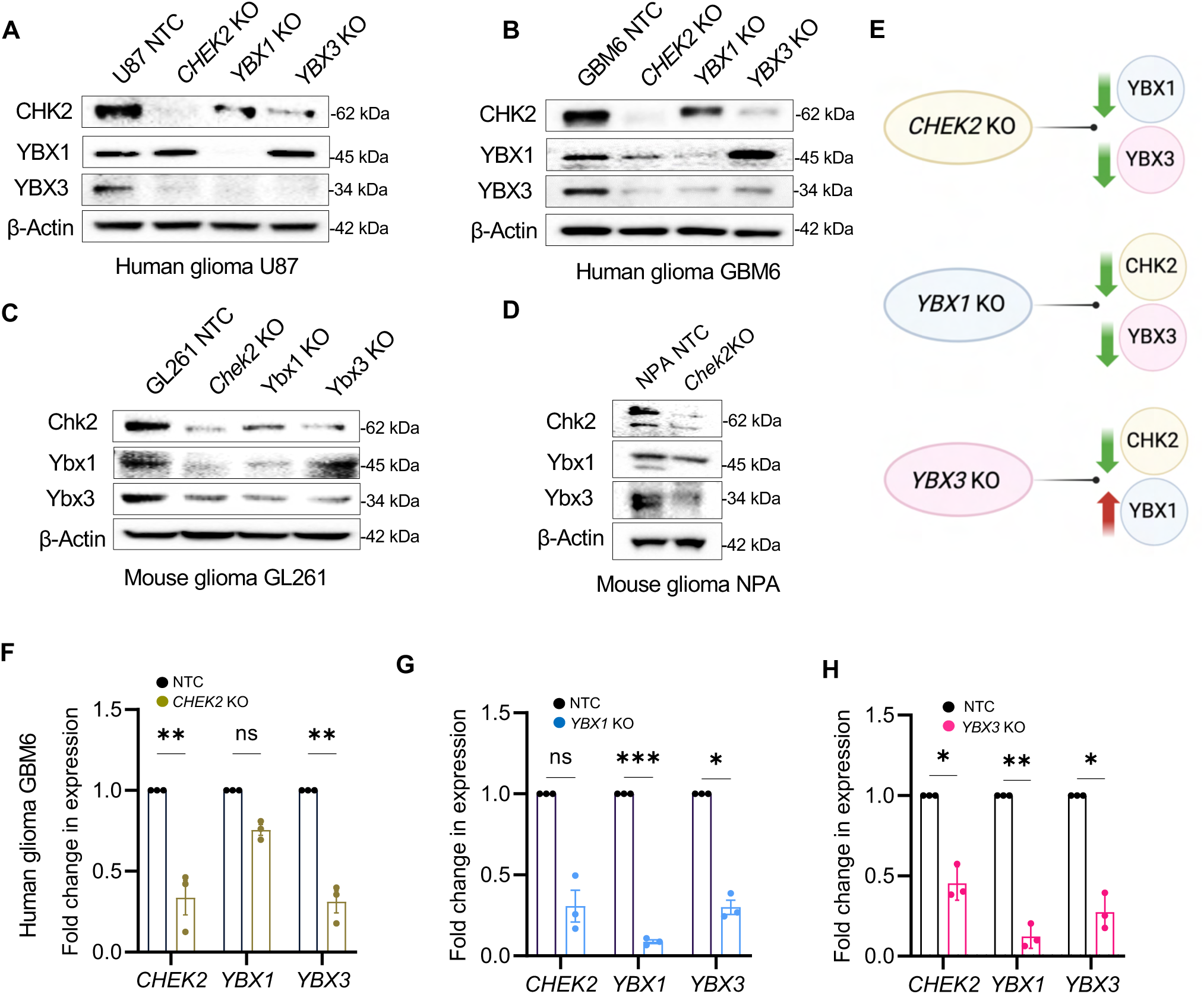
Reciprocal regulation between CHK2 and the YBX1/YBX3 hub at protein and transcript levels in GBM. (A) Immunoblots of single gene knockout of CHEK2, YBX1 or YBX3 in the human GBM cell line U87. The experiments were independently repeated three times. (B) Immunoblots of single gene knockout of CHEK2, YBX1 or YBX3 in the human GBM cell line GBM6. The experiments were independently repeated three times. (C) Immunoblots of single gene knockout of Chek2, Ybx1 or Ybx3 in the mouse glioma cell line GL261. The experiments were independently repeated at least three times. (D) Immunoblots of single gene knockout of Chek2 in the mouse glioma cell line NPA. (E) Schematic representation of gene expression changes in CHEK2 KO, YBX1 KO, and YBX3 KO conditions. Green downward arrows indicate decreased expression, while the red upward arrow represents increased expression. (F-H) RT-qPCR of CHEK2, YBX1 and YBX3 genes in human GBM6 cells with knockout of CHEK2 (F), YBX1 (G), and YBX3 (H). GAPDH was used to normalize target gene expression. The values were expressed as the fold change. Data are expressed as mean ± SE. N = 3 independent experiments with 3 technical replicates per experiment. Statistical differences between cell types were evaluated using one-way ANOVA.

To understand whether this regulation exist in murine glioma cells, we examined the mouse glioma cell lines GL261 and genetically engineered model derived cells called as NPA glioma cells (with genetic features (NRAS, shP53, shATRX, wt-IDH1)(31) that resembles human gliomas(32,33)). In GL261 cells, *CHEK2* KO led to a substantial reduction in both YBX1 and YBX3 protein levels, mirroring the regulatory patterns observed in GBM6 (Fig. 2C). *YBX1* KO further decreased CHEK2 and YBX3, whereas *YBX3* KO downregulated CHEK2 but induced a compensatory upregulation of YBX1. Likewise, in NPA cells, CHEK2 KO concurrently lowered YBX1 and YBX3 expression, reinforcing the functional conservation of this hub in murine glioma models (Fig. 2D).

To assess the transcriptional interdependence of the CHEK2-YBX1&YBX3 hub, reverse transcription-quantitative polymerase chain reaction (RT-qPCR) was performed in GBM6 following individual gene KO. In GBM6 cells (Fig. 2E–G), *CHEK2* KO (Fig. 2E) significantly reduced *CHEK2* and *YBX3* mRNA levels, indicating a strong regulatory link between these genes. *YBX1* KO (Fig. 2F) not only suppressed *YBX1* but also downregulated *YBX3*, whereas *YBX3* KO (Fig. 2G) led to a concurrent decrease in *CHEK2*, *YBX1*, and *YBX3* expression, highlighting a central role for *YBX3* in maintaining this regulatory hub. These results reveal a stronger reciprocal regulation in GBM6 and suggest that the CHEK2–YBX1&YBX3 hub exhibits context-dependent transcriptional control across different GBM models.

### CHK2-YBX1&YBX3 hub exerts transcriptional repression on pro-inflammatory programs

YBX1 and YBX3 are repressors of pro-inflammatory gene expression (34–36). To elucidate the role of CHK2, YBX1, and YBX3 in immune regulation within glioma cells, we performed bulk RNA sequencing (RNA-seq) analysis of U87 glioma cells, including non-target control (NTC), CHK2 KO, YBX1 KO, and YBX3 KO models (Fig. 3A-C). Differential gene expression analysis revealed extensive transcriptional reprogramming across all knockout conditions. Volcano plots highlighted widespread upregulation and downregulation of genes, underscoring the substantial impact of CHK2, YBX1, and YBX3 loss on the GBM transcriptome (Fig. 3A-C). Normalized to the non-targeting control with an adjusted P-value < 0.05, 3,285 genes were upregulated in *CHEK2* KO vs. NTC, 954 genes in YBX1 KO vs. NTC, and 1,326 genes in YBX3 KO vs. NTC. Conversely, 2,235 genes were downregulated in CHEK2 KO vs. NTC, 1,009 genes in YBX1 KO vs. NTC, and 696 genes in YBX3 KO vs. NTC.

**Figure 3:**
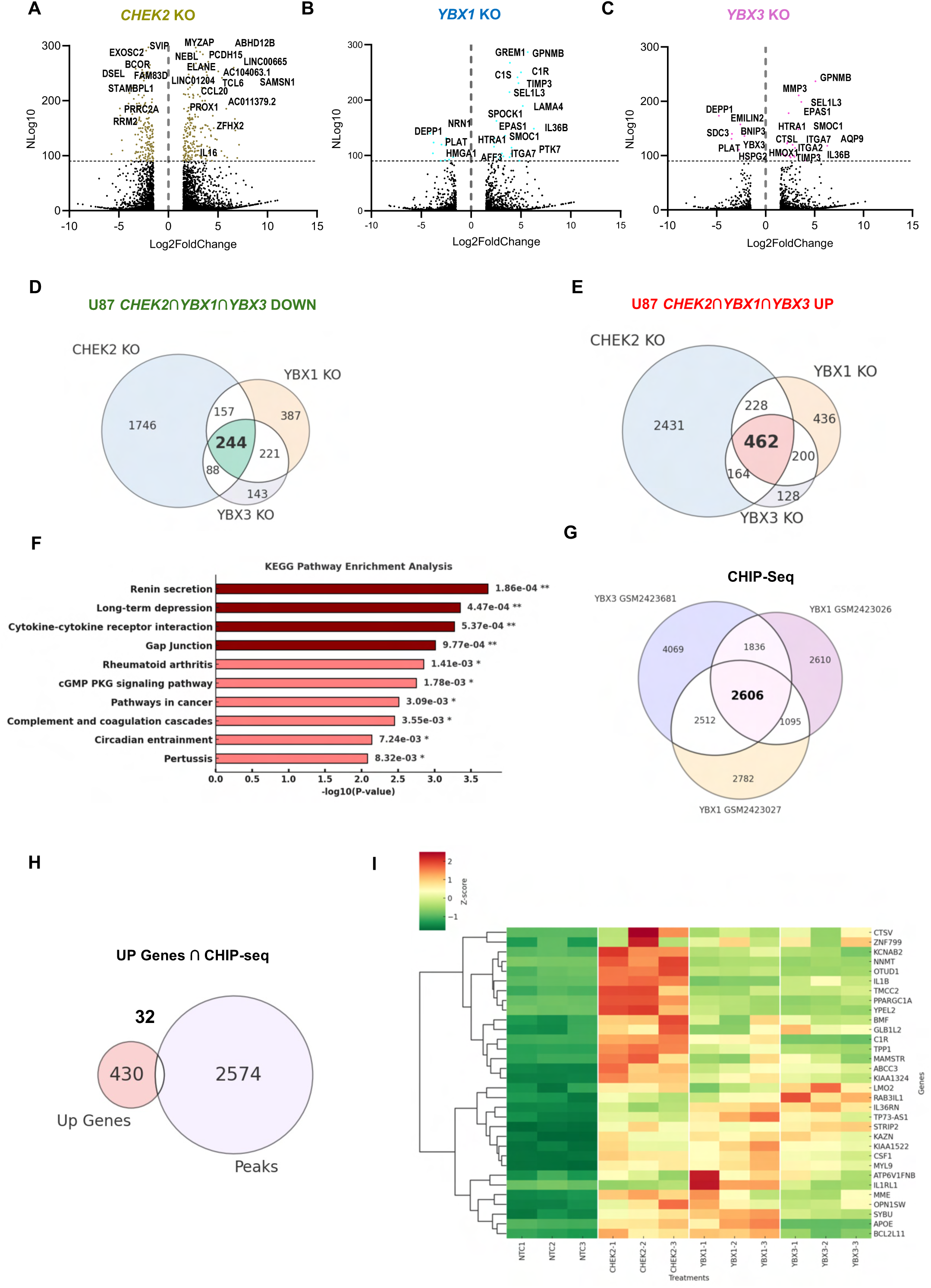
Genes regulated by CHK2-YBX1/YBX3 interaction: integration of RNA-seq and ChIP-seq data. (A) Volcano plot showing differentially expressed genes (upregulated and downregulated) in CHK2 knockout U87 cells. Upregulated genes are highlighted in the right half, and downregulated genes are in the left half. (B) Volcano plot showing differentially expressed genes in YBX1 knockout U87 cells. (C) Volcano plot showing differentially expressed genes in YBX3 knockout U87 cells. (D) Venn diagram illustrating the overlap of downregulated genes among CHK2, YBX1, and YBX3 knockout U87 cells, with the central red-labeled region indicating the 244 shared genes. (E) Venn diagram illustrating the overlap of upregulated genes among CHK2, YBX1, and YBX3 knockout U87 cells, with 462 shared genes in the center. (F) KEGG pathway enrichment analysis of the 462 overlapping upregulated genes, visualized as a bar graph. The x-axis represents the-log10(p-value), while the y-axis lists significantly enriched pathways. Darker green bars indicate more significant pathways. (G) Venn diagram showing ChIP-seq data from publicly available ENCODE datasets, illustrating 2,606 genes that are co-bound by YBX1 and YBX3. (H) Venn diagram depicting the overlap between ChIP-seq peaks (YBX1/YBX3-bound regions) and genes upregulated in CHK2, YBX1, and YBX3 knockout cells. The 32 identified genes are likely direct transcriptional targets. (I) Heatmap showing the expression of 32 upregulated genes across CHK2, YBX1, and YBX3 knockout conditions. The color scale represents Z-score normalized expression, where red indicates upregulation and green indicates downregulation. Hierarchical clustering was applied to group genes with similar expression patterns across knockout conditions.

To define shared transcriptional programs regulated by CHK2, YBX1, and YBX3, we examined overlapping differentially expressed genes (DEGs) across the three KO (Fig. 3D, E). A 462 number of genes were commonly dysregulated, suggesting a coordinated role for CHK2, YBX1, and YBX3 in transcriptional repression. Pathway enrichment analysis of these overlapping DEGs identified significant activation of inflammatory signaling, with cytokine-cytokine receptor interactions being one of the most enriched pathways (Fig. 3F). Several pro-inflammatory cytokines and receptors such as CCR1, IL1B, IL36B, CSF1, C1R and HLA-DQA1 were upregulated following CHK2, YBX1 and YBX3 individual KO cells, suggesting that this regulatory hub actively suppresses immune activation in GBM, Supplementary Fig. S1A.

To identify direct transcriptional targets of the CHK2-YBX1&YBX3 hub, we overlapped upregulated genes in CHK2, YBX1, and YBX3 KOs (Fig. 3E) with YBX1 and YBX3 chromatin immunoprecipitation sequencing (ChIP-seq) data (Fig. 3G) from the ENCODE project (37). Since YBX1 and YBX3 are transcriptional repressors, we selected the pool of 32 genes that were both bound by YBX1 and YBX3 in ChIP-seq dataset and were upregulated in all three conditions-*CHEK2*, *YBX1* and *YBX3* KO RNA seq dataset (Fig. 3H). Functional enrichment analysis revealed that these genes are associated with immune regulation and inflammatory response pathways (Fig. 3I). Similarly, we examined genes downregulated in CHK2, YBX1, and YBX3 KOs (Fig. 3D) and overlapped them with YBX1 and YBX3 ChIP-seq data (Fig. 3G). This analysis revealed 29 overlapping genes (Supplementary Fig. 2B, 2C).

Among these 32 upregulated genes, several function as key immune modulators. APOE, CSF1, CR1, and KIAA1522 regulate immune cell recruitment and antigen presentation, whereas IL1B, IL36RN, CTSV, and CSF contribute to cytokine and interferon-γ (IFN-γ) signaling (38–41). Additionally, genes such as BCL2L11 (BIM), MME (CD10), PPARGC1A, and ZNF799 play key roles in CD8⁺ T cell activation (42–45). The heatmap (Fig. 3I) illustrates the differential expression of these genes across knockout models, emphasizing their functional relevance in GBM immune suppression. This transcriptomic profiling suggests that targeting the CHK2-YBX1&YBX3 hub could upregulate a pro-inflammatory gene signature in glioma tumor cells.

### CHEK2-YBX1&YBX3 knockout gene signature aligns with the mesenchymal tumor cell state

Next, we assessed the relevance of our CHEK2-YBX1&YBX3 KO (Fig. 3 A-C) gene signature in the context of the human GBM landscape. To acquire a reliable reference for this expression pattern in human samples, we performed integrated analysis on publicly available single-cell (scRNA-seq) and spatial transcriptomics datasets (46) (Fig. 4A). Supplementary Fig. S2A shows the expression of CHEK2, YBX1 and YBX3 in these samples. The upregulated gene signature derived from CHK2, YBX1, and YBX3 KO U87 cells (Hub KO^high^) in malignant cell populations identified a distinct subset of tumor cells that was significantly enriched in mesenchymal (MES) and astrocyte (AC) like tumor cell subsets (chi-square test: p < 2,2×10^-16^; Fig. 4B-E; Supplementary Fig. S2B-D). Accordingly, hub KO^high^ cells showed selective enrichment of our RNA- and ChIP-seq overlap gene signature (Fig. 4F). We evaluated the spatial distribution of the newly identified hub KO^high^ positive tumor cells across our cohort of spatial transcriptomics samples, which showed a clear regional enrichment within the GBM microenvironment (Fig. 4G,H). Spatial correlation between cell types across our dataset showed an expected local association between KO^high^ positive tumor cells and MES-like/AC-like tumor cells (Fig. 4I). Importantly, KO^high^ positive tumor cells were most strongly correlated with naïve monocytes within the glioma TME. These results highlight that CHEK2-YBX1&YBX3 KO shift in-vitro tumor cell expression patterns towards a MES-like tumor state, a cell state that is known for their active engagement of the immune system through a host of mechanisms in the GBM TME, including MHC-I/II (47). Pathway enrichment analysis of hub KO^high^ regions consequently revealed a conserved association with epithelial-mesenchymal transition, extracellular matrix remodeling, immune activation, and interferon signaling (Fig. 4I, J). Notably, several immune-related pathways were shared between U87 RNA-seq and spatial transcriptomics, including cytokine-cytokine receptor interaction, TNF-alpha signaling, antigen presentation and processing, as well as complement and coagulation cascades, which are implicated in immune surveillance and tumor-immune interactions (Fig. 4I). Additionally, immune-related cancer pathways, including checkpoint regulation and inflammatory signaling, were commonly activated. Collectively, these findings establish a robust association between CHK2-YBX1&YBX3 signaling and immune modulation in human GBM tumors, demonstrating concordance between mechanistic insights derived from GBM cell line models and primary human GBM tumors.

**Figure 4.**
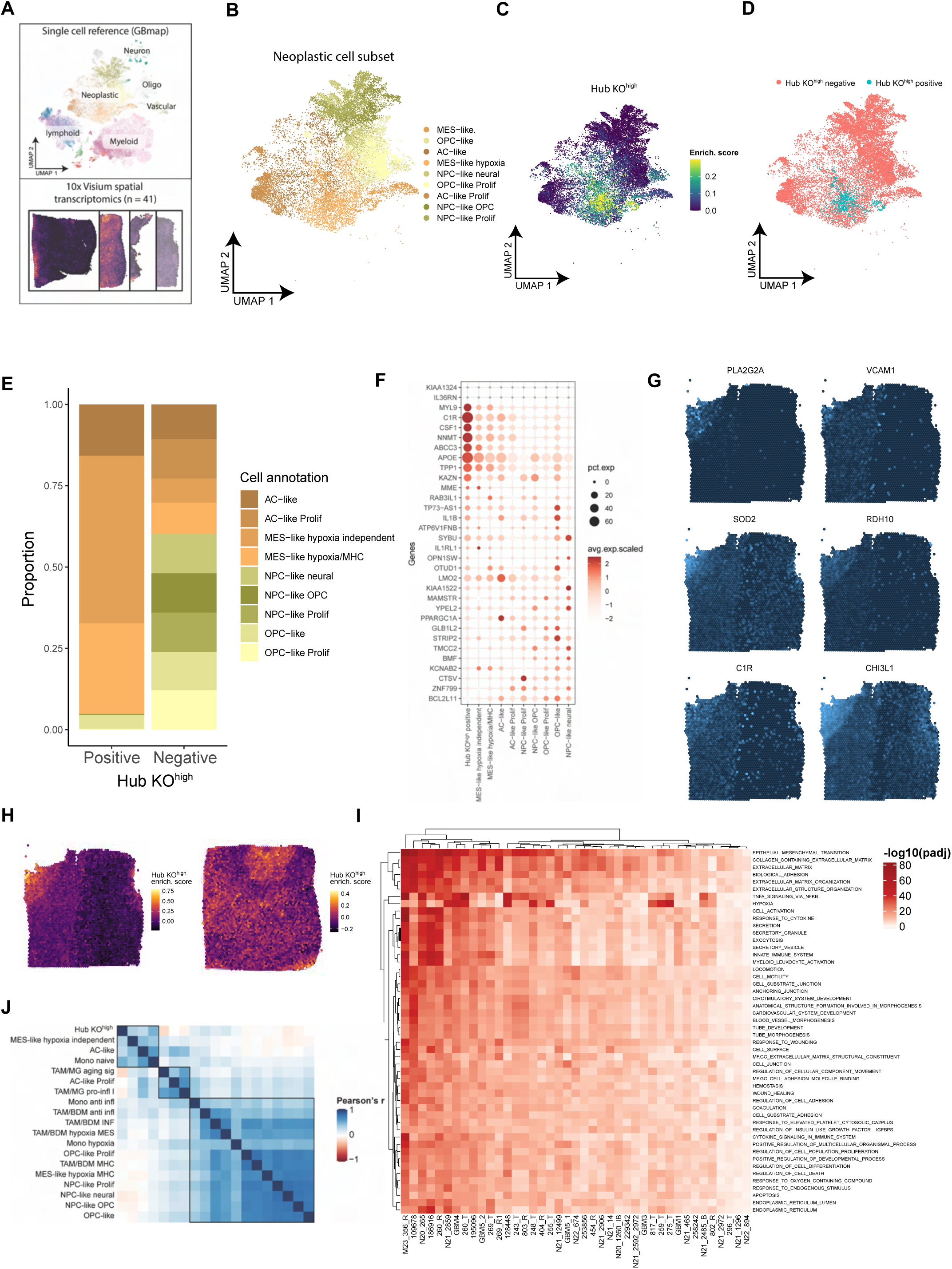
Spatial contextualization of CHEK2-YBX&YBX3 hub knockouts gene signature. (A) Overview of annotated GBM reference single cell RNA-seq and spatial transcriptomics data. (B) Neoplastic cell subpopulations of scRNA-seq reference in A. (C) Single cell enrichment scores for the consensus upregulated gene signature from Chek2, YBX1 and YBX2 KO of U87 human glioma. (D) Assignment of a Hub KO^high^ positive tumor cell subpopulation. (E) Distribution of the tumor subpopulations for Hub KO^high^ positive and negative tumor cells. (F) Scaled expression values of the genes identified from CHIP-Seq overlapping common upregulated genes of CHEK2, YBX1&YBX3 KO cells. (G) Spatial expression of marker genes from Hub KO^high^ positive tumor cell subcluster. (H) Two examples of spatial enrichment for the marker genes from the Hub KO^high^ positive tumor cell subcluster. (I) Spatial correlation between cell types in the tumor microenvironment and Hub Hub KO^high^ positive tumor cells. (J) Consensus of enriched gene sets associated with CHEK2-positive tumor cell regions across the spatial cohort.

### Repurposing the YBX1 inhibitor (SU056) to target the CHK2-YBX1&YBX3 regulatory hub in GBM

SU056, an azopodophyllotoxin small molecule, has been identified as a potent inhibitor of Y-box binding protein 1 (YBX1) (Fig. 5A) (29). Given that YBX1 is regulated by CHK2-YBX3 hub, we hypothesized that SU056 could disrupt this entire regulatory hub (Fig. 5B). While SU056 has been characterized for its ability to degrade YBX1, here we characterized its impact on YBX3 and CHK2. We evaluated the effects of SU056 in three GBM models: human U87 (Fig. 5C) and GBM6 (Fig. 5D) cells, as well as the murine GL261 cell line (Fig. 5E). Western blot analysis confirmed that SU056 treatment for 48 h led to YBX1 degradation, consistent with previous reports (29,48). Notably, SU056 also significantly reduced both YBX3 and CHK2 protein levels, marking a discovery of a yet unknown target. This degradation effect was consistently observed in both U87 and GBM6 cells, where CHK2 and YBX3 expression were markedly diminished following SU056 treatment. Furthermore, SU056 demonstrated pathway specificity by selectively targeting the CHK2-YBX1&YBX3 hub without affecting CHK2 related DNA damage response proteins ATM and CHK1.

**Figure 5:**
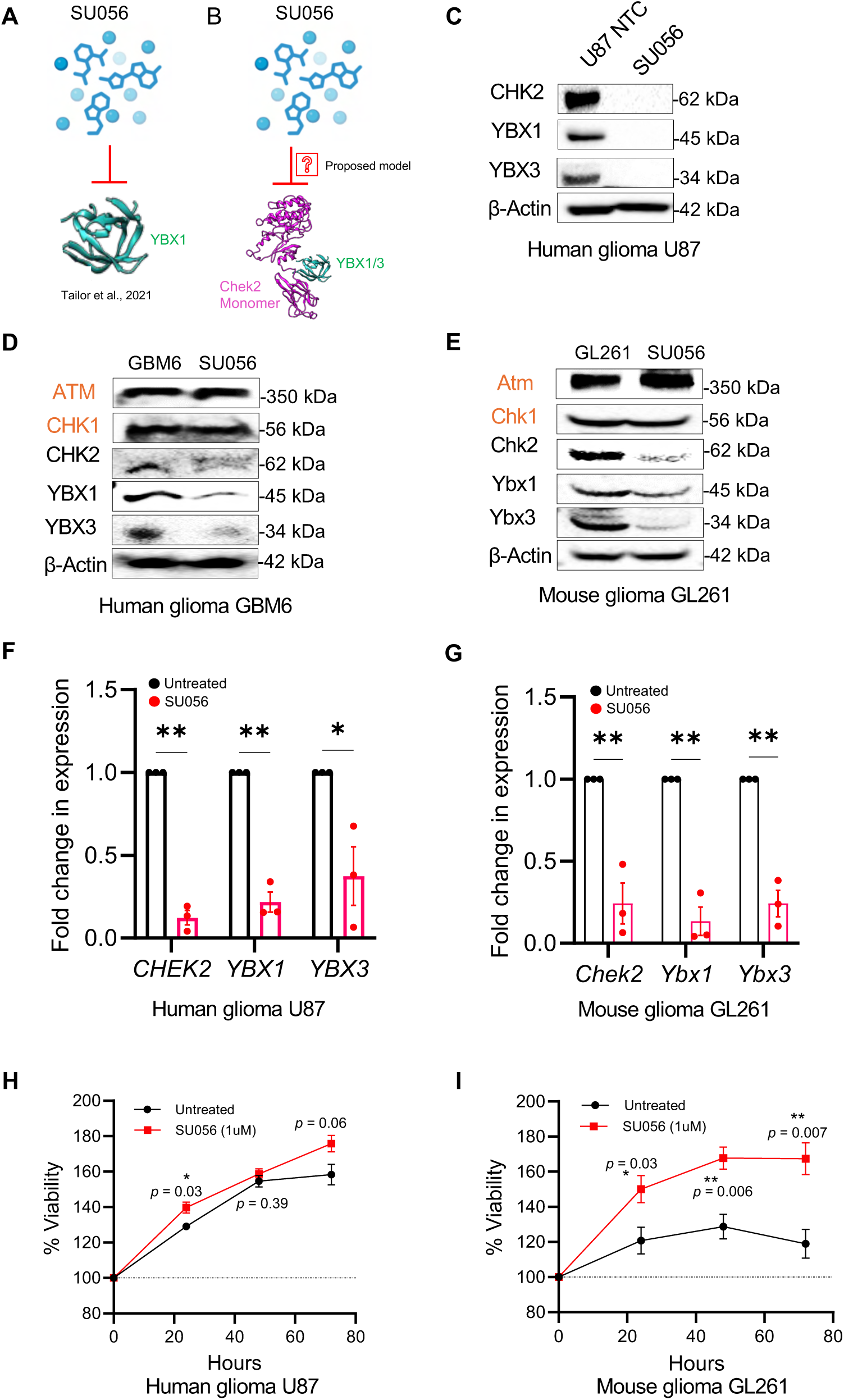
Targeting the CHK2-YBX1&YBX3 regulatory hub in GBM using the YBX1 inhibitor SU056. (A) Schematic representation of YBX1 inhibitor SU056 inhibiting YBX1 protein. (B) Proposed model of SU056 targeting the CHK2-YBX1&YBX3 hub. (C-E) Immunoblots of CHK2, YBX1, and YBX3 in human (C) U87 NTC (D) GBM6 and mouse (E) GL261 glioma cells treated with or without SU056 β-Actin. (F–G) RT-qPCR of CHK2, YBX1, and YBX3 mRNA expression in U87 (F) and GL261 (G) cells treated with or without SU056. The values were expressed as the fold change. Data are expressed as mean ± SE. N = 3 independent experiments with 3 technical replicates per experiment. (H–I) GBM cell viability assay. CellTiter-Glo® assay in U87 (H) and GL261 (I) cells following with or without SU056 treatment. N = 4 independent experiments.

To determine whether SU056 affects gene expression at the transcriptional level, RT-qPCR was performed to assess the mRNA levels of CHK2, YBX1, and YBX3 in human U87 and mouse GL261 cells following treatment. SU056 treatment for 48 h, resulted in a significant reduction in the transcript levels of all three genes across both models (Fig. 5F,G). Next, we assessed the impact of SU056 on GBM cell viability using the CellTiter Glo assay in U87 (Fig. 5H) and GL261 (Fig. 5I) cells. Cells were treated with SU056 (1 µM) for 24, 48, and 72 hours, and viability was measured relative to untreated controls. SU056 dose was not cytotoxic in U87 and GL261 cells. These results establishes that SU056 can deplete CHK2-YBX1&YBX3 levels.

### SU056 treatment enhances antigen presentation and CD8+ T cell expansion in glioma cells

YBX1, a known transcriptional repressor of immune-regulatory genes, has also been implicated in modulating type I and II interferon signaling (49–51). We use an unbiassed CellChat analysis to understand the communication of tumor cells with immune cells. The CellChat analysis identified MHC-I and MHC-II signaling to be significantly upregulated. Analysis of the MHC-I signaling network (Fig. 6A) revealed that hub KO^high^ tumor cell populations exhibited strong interactions with CD8^+^ T cells and NK cells relative to other tumor cell subpopulations. The MHC-II signaling network (Fig. 6B) demonstrated high interaction probabilities of hub KO^high^ tumor cells with plasma B cells and regulatory T cells (Tregs). Together, these results demonstrate that SU056 not only induces degradation of YBX1, YBX3, and CHK2 but also alleviates antigen presentation genes like MHC-I expression and enhanced CD8+ T cell activation. These findings support the CHK2-YBX1&YBX3 hub as a critical regulator of GBM immune evasion and position SU056 as a potential immune-modulatory agent for therapeutic intervention.

**Figure 6.**
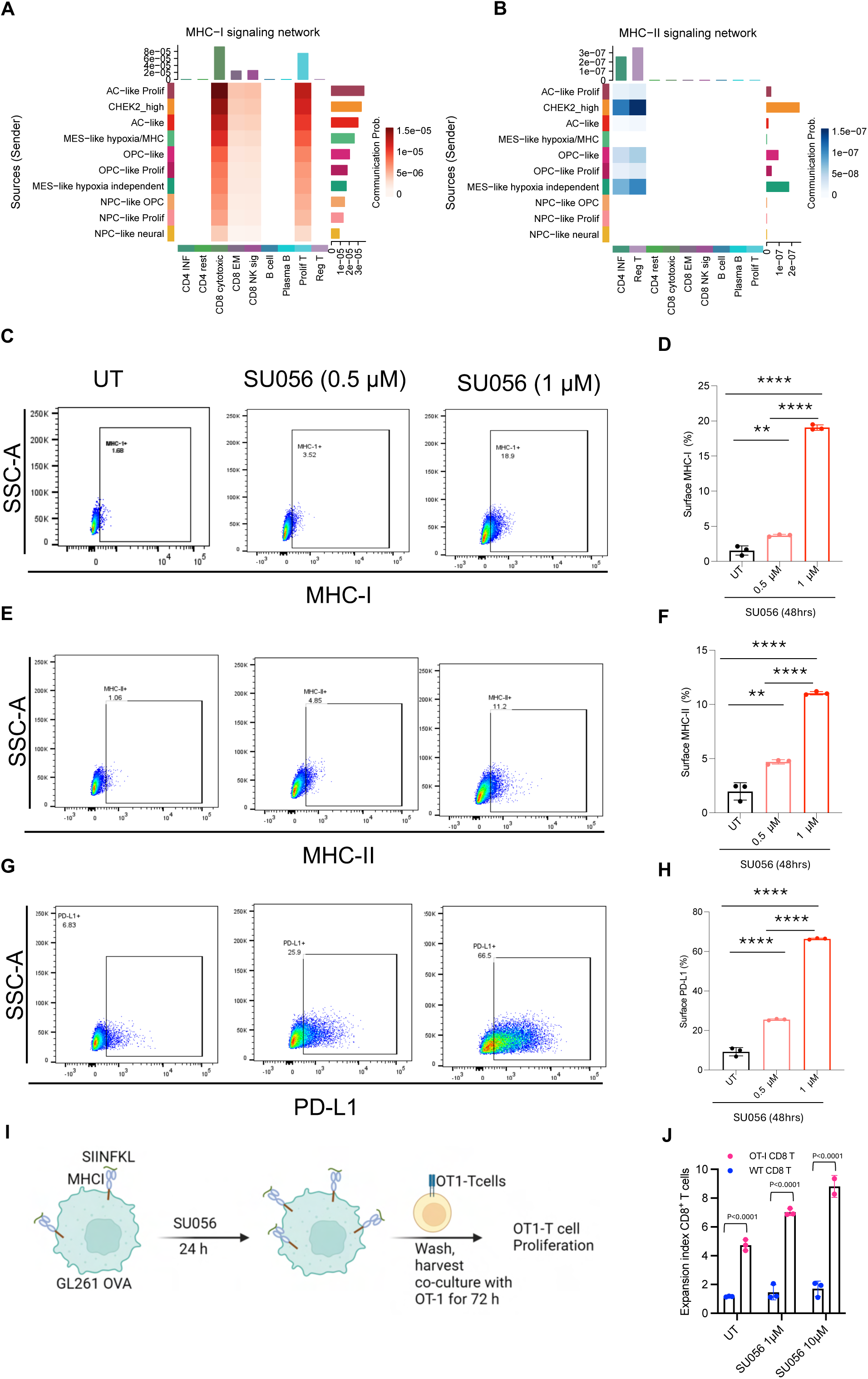
SU056 inhibition enhances IFNγ response, antigen presentation, and CD8+ T cell expansion. (A-B) CellChat analysis of scRNA-seq data showing interactions between tumor cell clusters (senders) and lymphocyte clusters (receivers) in the MHC-I (A) and MHC-II (B) signaling networks. (C-D) Flow cytometry representative and quantification of MHC class I expression on GL261 mouse GBM cells following 48-hour treatment with SU056 (0.5 µM and 1 µM), showing a dose-dependent effect. (E-F) Flow cytometry representative and quantification of MHC class II expression on GL261 cells after 48-hour SU056 treatment, showing a dose-dependent effect. (G-H) Flow cytometry representative and quantification of PD-L1 surface expression on GBM cells with or without SU056 treatment, indicating a dose-dependent expression. (I) Schematic illustrating the experimental workflow for assessing OT-I CD8+ T cell activation after SU056 treatment. GL261 ovalbumin (OVA) overexpressing cells were treated with SU056 for 24 hours, washed and co-cultured with OT-I T cells for 72 hours. (J) Quantification of CD8+ T cell expansion, showing the expression of T cell proliferation following SU056 treatment, indicating a dose-dependent expression.

To investigate this *in vitro*, GL261 mouse glioma cells were treated with increasing concentrations of SU056 (1 µM, 5 µM, and 10 µM), and antigen presentation was assessed via flow cytometry. SU056 induced a dose-dependent upregulation of MHC class I (Fig. 6C-D) and MHC class II (Fig. 6E-F) and PD-L1 expression (Fig. G-H). To functionally assess whether SU056-mediated immune modulation enhances anti-tumor immunity, GL261 overexpressing ovalbumin (OVA) cells pre-treated with SU056 and co-cultured with OT-I CD8+ T cells. SU056 treatment significantly increased CD8+ T cell expansion (Fig. 6I-J), further supporting its role in promoting antigen presentation and antigen specific CD8+ T cell proliferation.

Together, these results demonstrate that SU056 not only induces degradation of YBX1, YBX3, and CHK2 but also alleviates IFNγ response gene repression, leading to increased MHC-I expression and enhanced CD8+ T cell activation. These findings support the CHK2-YBX1&YBX3 hub as a critical regulator of GBM immune evasion and position SU056 as a potential immune-modulatory agent for therapeutic intervention.

### Combination therapy of SU056 and PD-1/PD-L1 blockade enhances survival in murine glioma models

To assess whether SU056 can target (deplete) CHK2-YBX1&3 levels in the brain of the glioma bearing mouse, we administered SU056 (20 mg/kg, intraperitoneally) to GL261 tumor–bearing mice for three consecutive days. Western blot analysis of the tumor containing brain tissue showed substantial degradation of YBX1, YBX3, and Chek2 proteins in SU056-treated tumors as compared to the vehicle-treated controls (Fig. 7B). Notably, immunohistochemical staining for CD8⁺ T cells revealed increased T-cell infiltration in SU056-treated tumors relative to vehicle controls (Fig. 7C-E).

**Figure 7:**
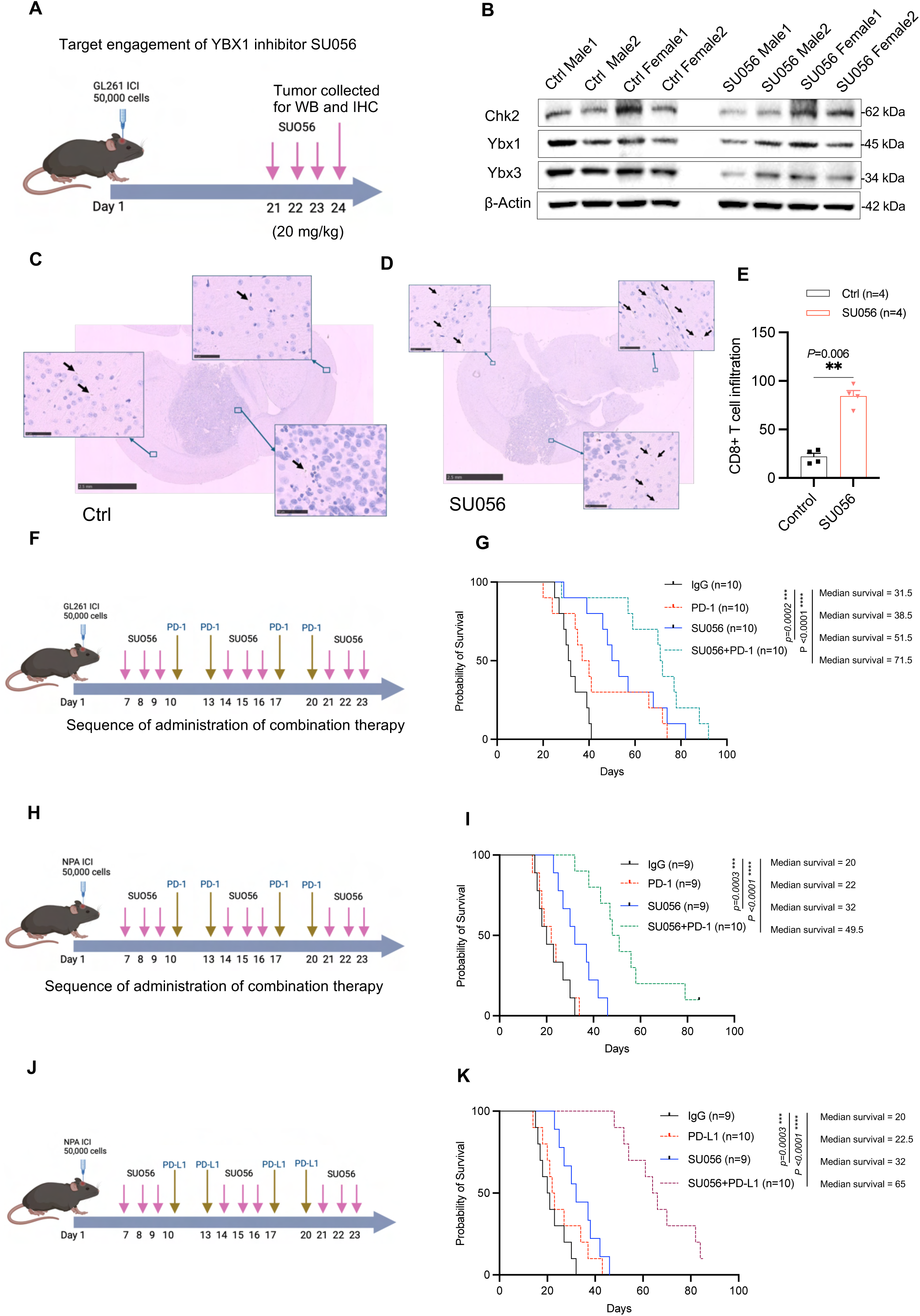
Therapeutic potential of YBX1 inhibition and PD-1/PDL-1 blockade in glioma. (A) Schematic representation of the experimental targeting engagement of SU056 timeline. C57BL/6 mice implanted intracranially with GL261 glioma cells, and administration of SUO56 started on day 21. Tumors were collected for WB and IHC analysis between on day 24. (B) Immunoblots of Chk2, Ybx1 and Ybx3 of tumor lysates harvested from brain tumor tissue of mice treated with SU056 or vehicle captisol. (C-D) Representative IHC staining of glioma sections for CD8a in vehicle-treated (C) and SU056-treated (D) mice. Black arrows indicate CD8a+ T cell infiltration into the tumor and brain microenvironment. Insets show magnified views of CD8a+ cells within the tumor. Scale bars: 50 µm, for small images. Scale bars: 2.5 mm, for full image. (E) Quantification of CD8a+ T cell infiltration in glioma sections from C. SU056 treatment significantly increased CD8a+ T cell infiltration compared with the control group. (**, P < 0.01, Student’s t-test). (F) Schematic of the treatment regimen for survival studies. Mice bearing intracranial GL261 gliomas treated with SU056 alone, PD-1 blockade alone, or the combination of SU056 and anti– PD-1 starting on day 7. (F) Kaplan–Meier survival curves for GL261 glioma-bearing mice. Mice were randomized into four groups (n = 10 per group): vehicle + IgG (VC + IgG), VC + anti–PD-1, SU056 + IgG, and SU056 + anti–PD-1. The median survival durations for each group were as follows: VC + IgG, 31.5 days; VC + anti–PD-1, 38.5 days; SU056 + IgG, 51.5 days; and SU056 + anti–PD-1, 71.5 days. Statistical comparisons: VC + IgG vs. SU056 + IgG (P =0.0002); and VC + IgG vs. SU056 + anti– PD-1 (P < 0.0001). (H) Schematic of the treatment regimen for survival studies in NPA glioma-bearing mice. Mice treated with SU056, PD-1 blockade, or their combination following tumor implantation. (I) Kaplan–Meier survival curves for NPA glioma-bearing mice treated as in G. Combination therapy with SU056 and PD-1 blockade significantly improved survival compared with single-agent treatments. Mice were randomized into four groups (n = 10 or n=9 per group): vehicle + IgG (VC + IgG), VC + anti–PD-1, SU056 + IgG, and SU056 + anti–PD-1. The median survival durations for each group were as follows: VC + IgG, 20 days; VC + anti–PD-1, 22 days; SU056 + IgG, 32 days; and SU056 + anti–PD-1, 49.5 days. Statistical comparisons: VC + IgG vs. SU056 + IgG (P =0.0003); and VC + IgG vs. SU056 + anti–PD-1 (P < 0.0001). (J) Schematic of the treatment regimen incorporating PD-L1 blockade in NPA glioma-bearing mice. Mice received SU056, PD-L1 blockade, or their combination. (K) Kaplan–Meier survival curves for NPA glioma-bearing mice treated. Mice were randomized into four groups (n = 10 or n=9 per group): vehicle + IgG (VC + IgG), VC + anti-PD-L1, SU056 + IgG, and SU056 + anti-PD-L1. The median survival durations for each group were as follows: VC + IgG, 20 days; VC + anti-PD-L1, 22 days; SU056 + IgG, 32 days; and SU056 + anti-PD-L1, 65 days. Statistical comparisons: VC + IgG vs. SU056 + IgG (P =0.0003); and VC + IgG vs. SU056 + anti-PD-L1 (P < 0.0001).

From the translational relevance standpoint, we next assessed whether SU056 could enhance the efficacy of immune checkpoint blockade in gliomas. Mice implanted with GL261 (Fig. 7F) and NPA glioma cells (Fig. 7H) were treated with SU056 monotherapy, anti-PD-1, or a combination of SU056 and anti-PD-1. Kaplan-Meier survival analysis demonstrated that SU056 monotherapy significantly extended median survival to approximately 51 days in GL261 and 32 days in NPA glioma models, compared to control mice. Notably, the combination of SU056 with anti-PD-1 further improved survival, extending median survival to 71 days in GL261 and 49 days in NPA, suggesting a therapeutic synergy between YBX1 inhibition and PD-1 blockade.

Since, SU056 treatment was found to upregulate PD-L1, we investigated whether combining SU056 with PD-L1 blockade could enhance therapeutic efficacy. To test this, we conducted a survival study using NPA tumor-bearing mice treated with SU056 in combination with anti-PD-L1 therapy (Fig. 7J). Like the PD-1 survival, SU056 monotherapy extended survival to 32 days, while in combination with PD-L1 blockade median survival increased to 65 days. These results establishes SU056 as a promising candidate for combinatorial immunotherapy strategies for gliomas.

## Discussion

Our findings identify a positive regulation between a DDR-related kinase and RNA/DNA binding proteins that is functionally implicated in this regulation is the repression of pro-inflammatory programs especially antigen presentation. Further, targeting of this regulatory hub using a single compound SU056 was found to be a therapeutically beneficial strategy for potentiating response to ICB in gliomas.

Single-gene KO experiments underscored that CHEK2, YBX1, and YBX3 form a regulatory hub, with each factor positively regulating the expression of the others. KO of any of the hub proteins led to downregulation of the expression of the other proteins at both protein and RNA levels. One exception is YBX3, as YBX3 KO led to the upregulation of YBX1 protein levels in both human and mouse glioma cells. This suggests that YBX3 KO may trigger compensatory stabilization of YBX1 protein levels as YBX3 KO significantly decreases YBX1 mRNA. Enhanced translation of YBX3 is known in the absence of YBX1, suggesting YBX1 negative control on YBX3 (52). Collectively, this suggest that YBX1 and YBX3 may substitute each other. However, the upregulated YBX1 levels in YBX3 KO cells were not able to restore the CHK2 protein levels, indicating that CHK2 and YBX3 regulation is non-compensable.

To investigate the functional consequence of the CHK2-YBX1&YBX3 regulation in gliomas, we performed the transcriptomic profiling of these cells. The upregulated gene repertoire from RNA-seq analysis of CHK2, YBX1 and YBX3 KO cells were involved in inflammatory signaling and immune activation including enhanced antigen presentation and cytokine interaction-related pathways. This indicated that CHK2-YBX1&YBX3 represses the pro-inflammatory transcriptional programs in glioma tumor cells to exert immunosuppression. Consequently, the loss of any one protein within this hub led to the de-repression of pro-inflammatory cytokines and immune-modulatory genes.

Building on these mechanistic insights, we explored the therapeutic potential of disrupting this hub using the YBX1 inhibitor SU056, which has previously been reported to degrade YBX1 in ovarian, pancreatic, and breast cancers (29,53,54). In glioma cell lines, SU056 not only depleted YBX1 but also led to the downregulation of CHK2 and YBX3, effectively degrading the CHK2-YBX1&YBX3 triad. This reprogramming was associated with alleviation of transcriptional repression on immune-stimulatory pathways including increased MHC class I and MHC class II expression and enhanced tumor-antigen presentation leading to enhanced CD8⁺ T cell expansion.

In our study, pharmacological inhibition of CHK2-YBX1&YBX3, rendered gliomas susceptible to ICB. Whereas we cannot rule out the contribution of depletion of other proteins, other than CHK2-YBX1&YBX3 by SU056, our mechanistic data using single gene KO of CHEK2, YBX1, and YBX3 indicate upregulation of proinflammatory gene signature including antigen presentation(17,22,29,48,52,53,55).

Our study has several limitations. Although we have identified the CHK2-YBX1&YBX3 hub, there is a possibility of other proteins being part of this hub/regulation. Similarly, we have investigated the effect of SU056 on the degradation of CHK2-YBX1&YBX3 hub. However, SU056 may have other unknown targets that could play a role in the overall effect of the SU056 treatment. While we have used multiple murine glioma models for the initial testing of the combination therapy, they may fail to fully recapitulate the biology of human GBMs and hence response to the therapy. Moreover, the exact binding sites involved in the interaction between CHK2-YBX1&YBX3 sites need to be studied using structural biology approaches.

In summary, we identified a CHK2-YBX1&YBX3 hub that exerts immunosuppression by transcriptionally repressing the pro-inflammatory networks in glioma tumor cells. Depletion of the CHK2-YBX1&YBX3 hub by SU056 compound restored the pro-inflammatory pathways and enhanced response to ICB in murine glioma models. Since CHK2-YBX1&YBX3 deletion is associated with enhanced antigen presentation accompanied by CD8 T cell proliferation, this therapy has the potential to be combined with alterative T-cell based immunotherapies for immunotherapy resistant cancers. To conclude, this work proposes a rationale and preclinical evidence to target CHK2-YBX1&YBX3 hub with ICB in gliomas.

## Methods

### Mouse models

All mice were housed at the Center for Comparative Medicine at Northwestern Feinberg School of Medicine. Mice were housed in a conventional barrier facility with 12-h light/12-h dark cycles and ad libitum access to food and water. C57BL/6 mice were obtained from The Jackson Laboratory. All experiments were performed on 6–8 weeks old mice, age- and gender-matched. For all survival studies, C57BL/6 mice were between 6–8 weeks old, and numbers of male/female mice were always equivalent between control and experimental groups. All mouse protocols performed in this study were approved by Northwestern’s Institutional Animal Care and Use Committee (IACUC) under study approval number IS00028126.

### Cell lines and tumor implantation

HEK293T cells were obtained from ATCC (Catalog no. CRL-3216), while the GL261 mouse glioma cell line was sourced from the National Cancer Institute (NCI). Both cell lines were cultured in Dulbecco’s modified Eagle’s medium (Corning) supplemented with 10% fetal bovine serum (FBS; HyClone) and penicillin-streptomycin. Additionally, NPA cells were kindly gifted by Prof. Maria Castro (University of Michigan). These cells are cultured as neurospheres in a medium composed of DMEM/F12, supplemented with 1× B27, 1× N2, 1× Normocin, and 1× antibiotic/antimycotic, along with human recombinant EGF and basic-FGF at concentrations of 20 ng/mL each. GBM6 patient derived xenograft (PDX) cells purchased from Mayo Clinic (Scottsdale, AZ, USA) and U87 from ATCC (Catalog no. HTB-14) were grown in Dulbecco’s Minimum Essential Media (DMEM) containing 10% FBS. The GBM6, U87, HEK293T, GL261, and NPA cell lines used in this study were authenticated annually via short tandem repeat profiling and were tested for mycoplasma contamination on a semi-annual basis. All assays were conducted using cell lines that were both STR-verified and free from mycoplasma. All cell lines were authenticated and regularly tested for mycoplasma contamination and maintained at 37°C in a 5% CO₂ incubator.

### Intracranial Immunocompetent Mouse Model

The intracranial implantation and monitoring protocol IS00028126 was approved by the IACUC at Northwestern University. Wild-type C57BL/6 mice were maintained in a pathogen-free environment at a stable 24 °C with humidity between 30–50%. Male and female mice aged 6–8 weeks were used, with wild-type animals (B6.129S2-Cd8atm1Mak/J; Strain #:002665) obtained from Jackson Laboratory.

For generating the intracranial models, mice were first anesthetized via an intraperitoneal injection of a ketamine (100 mg/kg) and xylazine (10 mg/kg) mixture. After disinfecting the surgical area, a midline incision was made to access the skull. Then, 5 × 10^4 GL261 or NPA cells were implanted to induce orthotopic tumors. Specifically, a 2.5 µl cell suspension in sterile PBS was loaded into a 29 G Hamilton syringe and slowly injected over three minutes into the left hemisphere at a 3 mm depth, through a transcranial burr hole created 3 mm lateral and 2 mm caudal to the bregma. Once the implantation was complete, the incision was closed with 9 mm stainless steel wound clips, and the mice were placed in a clean cage on a heating pad until recovery.

Throughout the study, the animals were monitored daily, and euthanasia was performed when they reached endpoint criteria (defined as >20% weight loss, loss of mobility, or severe neurological impairments such as seizures or circular motion) as specified in the IACUC protocol. Additionally, a balanced 1:1 male-to-female ratio was maintained during treatment randomization, and the tumor injection site and depth were consistently preserved across all experiments.

### Immunoblotting

Cells were lysed in RIPA buffer (Thermo Scientific, #89900) supplemented with a Protease Inhibitor Cocktail (Thermo Scientific, #78429). Protein concentration was determined using the BCA Assay (Thermo Scientific, #PI23225). Equal amounts of protein were separated by SDS-PAGE and transferred onto nitrocellulose membranes (Bio-Rad, #1620112). Membranes were incubated overnight at 4°C with primary antibodies: CHK2 (#2662S), YBX1 (#4202S), β-Actin (#4970S), ATM (#2873S), and CHK1 (#2360S) from Cell Signaling, and YBX3 (#TA324558) from Origene. HRP-conjugated secondary antibodies (anti-mouse, #7076S; anti-rabbit, #7074S, Cell Signaling) were applied, and detection was performed using an ECL substrate with the ChemiDoc Imaging System (Bio-Rad). Results are based on three independent experiments, each with three technical replicates.

### Generation of knockout clones

The guides for YBX1, YBX3 and NTC were cloned in lentiCRISPR v2-Blast vector ( The vector backbone was purchased from Addgene-LentiCRISPR v2-Blast was a gift from Mohan Babu (Addgene plasmid # 83480; http://n2t.net/addgene:83480; RRID:Addgene_83480). The protocol for guide cloning and generation of the virus was as described in (56). The U87, GBM6 and GL261 YBX1, YBX3 and NTC control clones were selected and maintained in Blasticidin (5 µg/ml). The mouse Chek2 KO and control plasmids described in our previous study (17) were used to generate Chek2 KO in mouse glioma GL261 and NPA lines. The selection conditions were: GL261 (2 µg/ml puromycin) and NPA (1 µg/ml puromycin). The plasmid for human Chek2 KO (CHEK2 C3.2 gRNA) was a gift from Iain Cheeseman (Addgene plasmid # 90628; http://n2t.net/addgene:90628; RRID:Addgene_90628) (57). The human Chek2 KO plasmid was transduced in a cas9 expressing GBM6 and U87 cells. The guide sequences for Chek2, YBX1 and YBX3 and NTC are shown in Supplementary Table1. For the purpose of ovalbumin overexpression, the pAc-Neo-OVA plasmid was acquired from Addgene (58). The GL261 cells were transfected with this plasmid and subsequently selected and maintained in G-418 (200 µg/ml; Sigma-Aldrich).

### Quantitative real-time PCR

Quantitative real-time PCR (qPCR) was then performed on a Bio-Rad real-time PCR system using the iTaq Universal SYBR Green Supermix (Bio-Rad, Cat. #1708880). Relative gene expression was determined by the ΔΔCᵀ method, with GAPDH as the internal control. Primer sequences for target genes, including CHEK2, YBX1, and YBX3, are listed in Supplementary Table 2. Data are expressed as mean ± SE. Results are based on three independent experiments, each with three technical replicates.

### Single cell and spatial transcriptomics data

We used a large, publicly available, well annotated single cell RNA-sequencing (scRNA-seq) reference data containing GBM samples from 110 patients (46). We also expanded the scRNA-seq data to include single cell profiles of neurons. Data acquisition for the 41 spatial transcriptomics samples that were used for the analysis was described previously (59,60). An overview of the patient information can be found in supplementary materials.

### Single-cell deconvolution using Cell2location

To evaluate cell type locations in the Visium spatial transcriptomics data, we used the Cell2location model with the abovementioned scRNA-seq data as a reference. We setup the Cell2location model by configuring an AnnData object, defining parameters such as the number of cells per location and detection sensitivity. The model was trained on a graphics processing unit for 500 iterations. After training, we extracted the posterior distribution of cell type proportions, sampling 1,000 times for precise estimation across the spatial framework. Median estimates of cell type abundance were saved in the AnnData object. Finally, we incorporated the cell type abundance data back into the SPATA object using the addFeature function in SPATA2, allowing for further analysis steps within the spatial data framework (61).

### Enrichment scores and cell annotation of gene signatures in single cell and spatial transcriptomics

Enrichment scores for CHEK2-YBX1&YBX3 knockout gene signatures in scRNA-seq and spatial transcriptomics data were calculated as follows: for a gene set G a reference gene set R was constructed. This was done by binning the all genes from the expression dataset into 30 equal-sized bins according to expression level. For every gene in G, 100 random genes from the corresponding bin were added to R, resulting in two gene sets, so that R was 100x larger than G. The difference in average expression for G and R was consequently calculated for each cell to provide an enrichment score. Gene sets that were used for enrichment calculations were filtered based on the top 2000 most variable genes in the scRNA-seq dataset. Thresholds for enrichment scores was defined as the mean plus one standard deviation of the enrichment scores across all cells. Differentially expressed genes for cell clusters in snRNA-seq data were determined using the *FindMarkers* function from the Seurat R package. For hub KO^high^ positive tumor cells, marker genes used in spatial transcriptomics analysis were selected using log2 fold change cutoff of larger than 3 and an adjusted p value of smaller than 0.01.

### Spatial transtricptomics correlation and gene set enrichment analysis

Spatial enrichment scores for the hub KO^high^ positive tumor cell signature was calculated for individual Visium spots as described above. From our cohort of spatial transcriptomics GBM samples, we selected samples with a standard deviation of over 0.08 for the KO^high^ signature to restrict the analysis to samples with meaningful spatial enrichment, which resulted in 42 samples. Deconvoluted cell type abundance from the Cell2location algorithm were used for cell type specific correlation analysis with spatial enrichment scores for the KO^high^ signature using Pearson’s r to determine correlation. Gene set enrichment analysis was performed using the SPATA2 package with all gene sets included in standard workflow. The 50 gene sets that showed the highest average significance across all samples were selected.

### Ligand-receptor interaction analysis

Cell-cell interaction inference was performed in our integrated snRNA-seq dataset using the CellChat.DB R package. We included all standard cell-cell interaction pathways that are included in the CellChat.DB package. Standard inference analysis was used with the default 25% truncated mean (triMean) setting (60). Centrality scores for annotated cell types are a measure of importance in a specific pathway as a “sender” or “receiver” and was adopted from the standard CellChat.DB output.

### *In vitro* antigen presentation and T-cell proliferation assays

For ovalbumin overexpression in GL261, the pAc-Neo-OVA plasmid was purchased from Addgene (62). GL261 cells were transfected with the pAc-Neo-OVA plasmid and were selected and maintained in G-418 (200 µg/ml; Sigma-Aldrich).GL261 mouse glioma cells stably expressing ovalbumin (GL261-OVA) were allowed to adhere overnight. The following day, cells were treated with SU056 or vehicle control for 24 hours. After treatment, cells were gently detached using EDTA (to preserve surface epitopes) and stained to assess surface expression of MHC-I (clone AF6-88.5.5.3, BIB515, eBiosciences), MHC-II (clone AF6-120.1, BV605, Invitrogen) and PD-L1 (clone 10F.9G2, PE-Cy7, BioLegend) at a 1:100 dilution. Data were acquired on a BD Symphony flow cytometer and analyzed with FlowJo (version 10.10.0).

For T-cell proliferation assays, CD8⁺ OT-1 T cells were isolated from spleens of OT-1 transgenic mice using a commercial T-cell isolation kit (e.g., Mouse T-cell Isolation Kit, StemCell Technologies, Cat. No. 19853). These T cells were labeled with eBioscience Cell Proliferation Dye eFluor 450 (5 μM, Thermo Fisher). Labeled OT-1 T cells were then co-cultured with pretreated GL261-OVA cells at a 5:1 T cell–to–tumor cell ratio for 72 hours in complete RPMI medium. After 72 hours, T cells were harvested, blocked with Fc-blocking reagent (anti-CD16/32), and stained for CD8 (e.g., CD8a BV605, clone 53-6.7, BioLegend) and viability (Fixable Viability Dye eFluor 780). Proliferation was assessed by dye dilution on a BD Symphony flow cytometer, and data were analyzed using FlowJo (version 10.7.1). Data are expressed as mean ± SE. Results are based on three independent experiments, each with three technical replicates.

### Therapy and dosing for mouse survival studies

Orthotopic glioma models were generated by intracranially implanting 5×10^4 GL261 or NPA cells into immunocompetent mice. Beginning on day 7 after tumor implantation, animals were randomized to treatment groups according to the schedules illustrated in the figure panels. Treatment arms included vehicle control, SU056 monotherapy (20 mg/kg, administered intraperitoneally on specified days), anti-PD-1/anti-PD-L1 (200 µg/dose, injected intraperitoneally as indicated), and various combination therapies incorporating SU056 with either anti-PD-1 or anti-PD-L1 antibodies (200 µg/dose, Bio X Cell). On days when anti-PD-1 or anti-PD-L1 was administered, an isotype control antibody (200 µg/dose, rat IgG2a, Bio X Cell) was administered in the control cohorts. The treatment was stopped beyond day 23 post intracranial injection. Kaplan–Meier survival analysis was conducted to determine therapeutic efficacy.

### Drug preparation for in vivo dosing

SU056 was initially dissolved in DMSO and then diluted into 20% Captisol (2 g Captisol dissolved in 10 mL of 0.9% saline) to achieve the required working concentration. This solution was stored at 4°C and used within 14 days. Animals in the vehicle-control groups received either 20% Captisol or PBS following the same schedule and injection volume as the SU056-treated groups.

### Immunohistochemistry for CD8⁺ T-cell infiltration

Four tumor-bearing mouse brains per group were collected at the experimental endpoint and transferred to the Mouse Histology and Phenotyping Laboratory at Northwestern University for tissue processing. Formalin-fixed, paraffin-embedded (FFPE) tumor specimens were cut into 5 μm sections, deparaffinized in xylene, and rehydrated in a series of graded alcohol solutions. Antigen retrieval was performed in sodium citrate buffer (pH 6.0) at 95°C for 20 minutes. Endogenous peroxidase activity was quenched using 3% hydrogen peroxide, and non-specific binding was blocked with 5% normal serum in PBS for 30 minutes at room temperature.

Sections were then stained with a mouse anti-CD8 antibody (Cell Signaling Technology cat # 98941S) at a 1:200 dilution on a DAKO Autostainer Link 48 (Agilent Technologies). After washing in PBS, slides were incubated with the appropriate secondary antibody, followed by visualization with a peroxidase-based detection system and 3,3′-diaminobenzidine (DAB) substrate. Adjacent sections were stained with hematoxylin and eosin (H&E) for histopathologic evaluation. Finally, slides were counterstained with hematoxylin, dehydrated, and mounted with coverslips. All sections were scanned using a Hamamatsu NanoZoomer 2.0 HT, and images were reviewed in NDP.view2 software. Isotype IgG–treated sections served as negative controls. CD8⁺ T cell IHC quantification was performed in a blinded manner by counting all stained cells across the entire brain section, including both tumor and non-tumor areas in NDP.view2 softwaare. Four brains from the control group and four brains from the SU056-treated group were analyzed. Data are expressed as mean ± SE. Results are based on three independent experiments, each with three technical replicates.

### RNA sequencing

*CHEK2* knockout, *YBX1* KO, *YBX3* KO, and non-targeting control (NTC) glioma cells were cultured under standard conditions and harvested at 70–80% confluence. Total RNA was extracted using Zymo RNA isolation kit, in accordance with the manufacturer’s protocol. RNA integrity was then evaluated on an Agilent 2100 Bioanalyzer, and only samples with an RNA integrity number (RIN) ≥7 were used to construct sequencing libraries.

Library preparation was performed using a conventional mRNA workflow, which involved poly(A) selection, cDNA synthesis, adapter ligation, and PCR amplification. All RNA-seq libraries were processed and sequenced at Novogene on an Illumina platform, producing 150 bp paired-end reads. Data generated from these experiments were used to investigate the transcriptomic changes associated with the CHK2-YBX1&YBX3 regulatory axis in glioma cells. Data are expressed as mean ± SE. Results are based on three independent experiments, each with three technical replicates.

### Statistical analyses

Flow cytometry data was analyzed by FlowJo™ v10.10 software. Statistical analyses were performed using Graphpad Software (Prism v10.04). Student’s t-test was used to measure statistical differences between two groups. One-way or two-way analysis of variance was used for multiple comparisons, and p values were adjusted for multiple comparisons where appropriate. Survival curves were generated via the Kaplan–Meier method and compared by the log-rank test. All the tests were two-sided and p values less than 0.05 were considered significant.

## Data Availability

The raw data related to RNA sequencing is available on the publicly available repository, Sequence Read Archive (SRA) Submission ID: SUB15161080, under accession number PRJNA1233302 (https://www.ncbi.nlm.nih.gov/bioproject/PRJNA1233302). The publicly available ChIP-seq data used in this study in available as GSM2423026, GSM2423027 and GSM2423681(63). The schematic diagrams presented in this study are made using BioRender (www.biorender.com). The authors have the biorender license that permits content to be sublicensed for use in journal publications. All data supporting the findings of this study are available within the article and its supplementary materials information files. Source data are provided with this paper.

## Competing interest statement

No potential conflicts of interest were disclosed.

## Acknowledgments

This work was supported by R01NS138769 to CD, NIH/NCI P50CA221747 SPORE for Translational Approaches to Brain Cancer (Dr. Maciej Lesniak) and 5DP5OD021356, 1R01CA245969 to AMS and R01NS096376-06A1 to AUA. We thank the Mouse Histology & Phenotyping Laboratory and the Pathology Core Facility Laboratories at Northwestern University for performing immunohistochemistry and imaging of the whole slides.

---

**Supplementary Figure S1.**
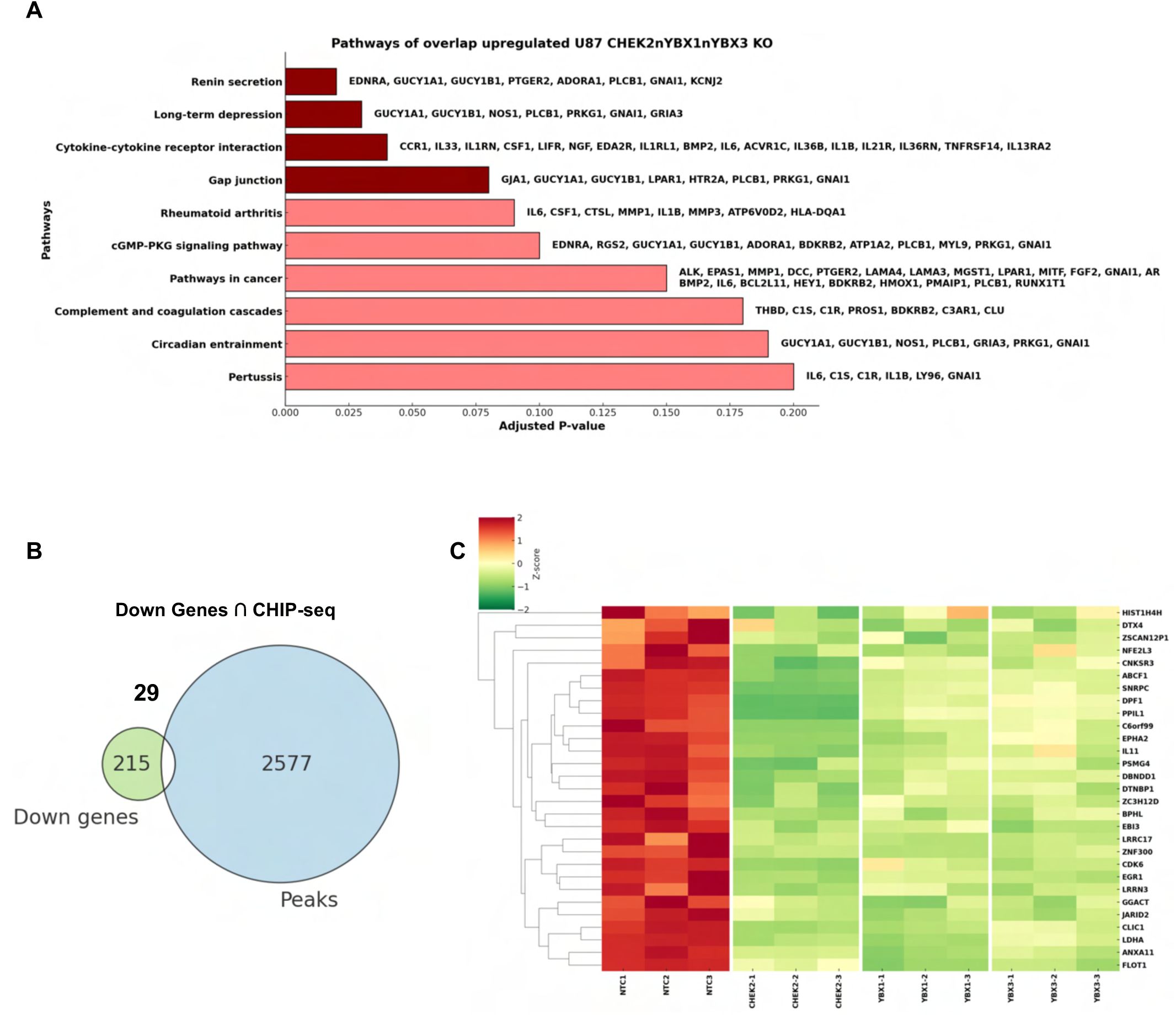
(A) Pathway enrichment analysis of upregulated genes overlapping in CHK2, YBX1, and YBX3 knockout (KO) U87 cells. The bar graph represents significant pathways, showing the adjusted p-values along the x-axis and the associated genes within each pathway. (B) Venn diagram illustrating the overlap between downregulated genes at the RNA level (from CHK2, YBX1, and YBX3 KO cells) and genes transcriptionally regulated by YBX1 and YBX3 (identified through ChIP-seq). A total of 29 genes were identified as common between the two datasets. (C) Heatmap representing the expression levels of the 29 downregulated genes in CHK2, YBX1, and YBX3 KO cells. The color gradient represents the Z-score normalized expression, with red indicating upregulation and green indicating downregulation. Hierarchical clustering was applied to visualize gene expression patterns across different KO conditions.

**Supplementary Figure S2.**
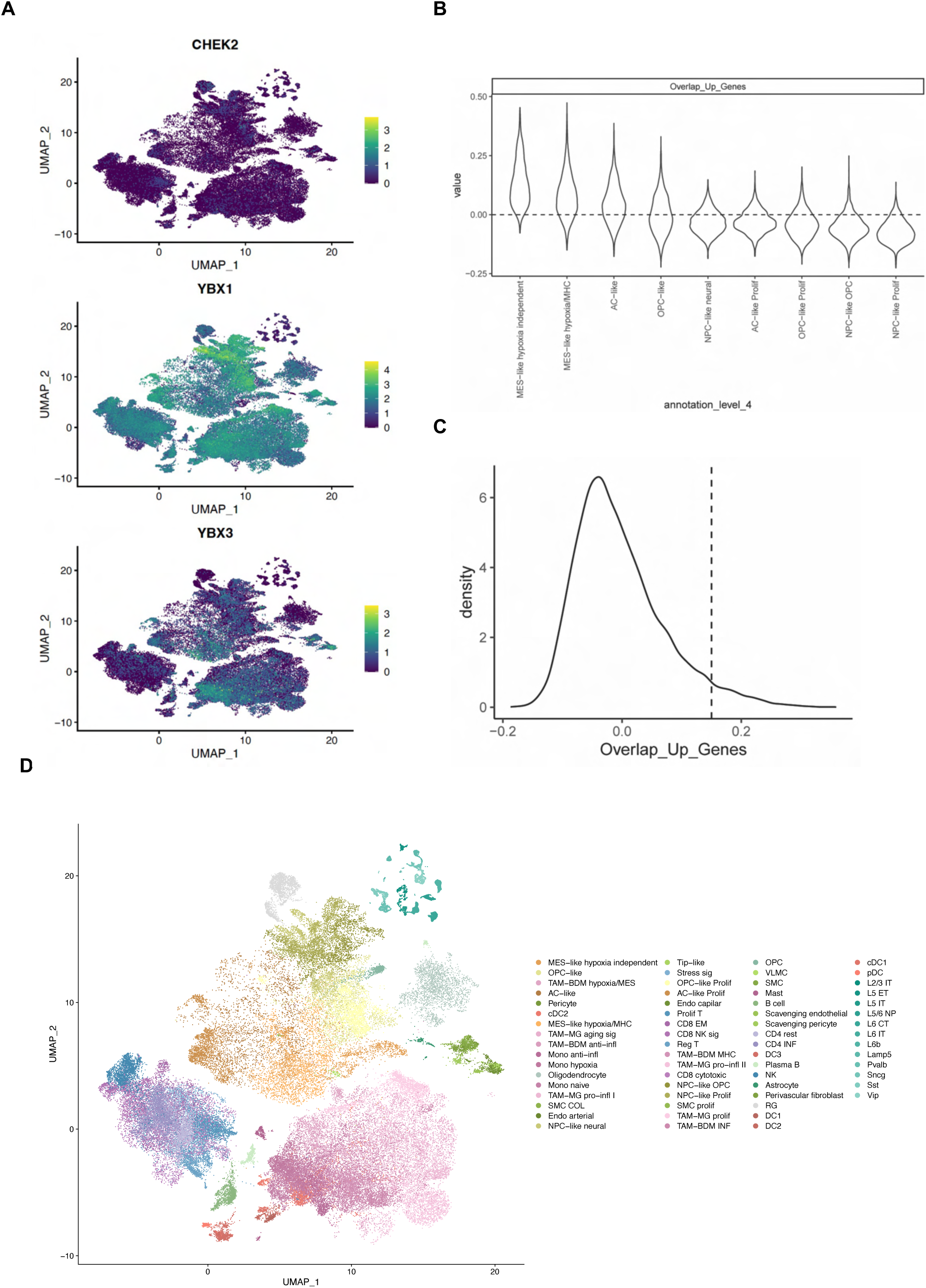
(A) Single-cell RNA sequencing UMAP plots showing the expression of CHEK2, YBX1, and YBX3 across different cell clusters. The color scale indicates the expression levels of each gene, where purple represents low or no expression, and yellow-green represents high expression. (B) Enrichment score distribution in all tumor cell subsets. (C) Density plot showing the cutoff for Chek2 gene set enrichment score in tumor cells. (D) Annotation of complete scRNA-seq reference including Hub KO^high^ positive tumor cells.

**Supplementary Figure S3.**
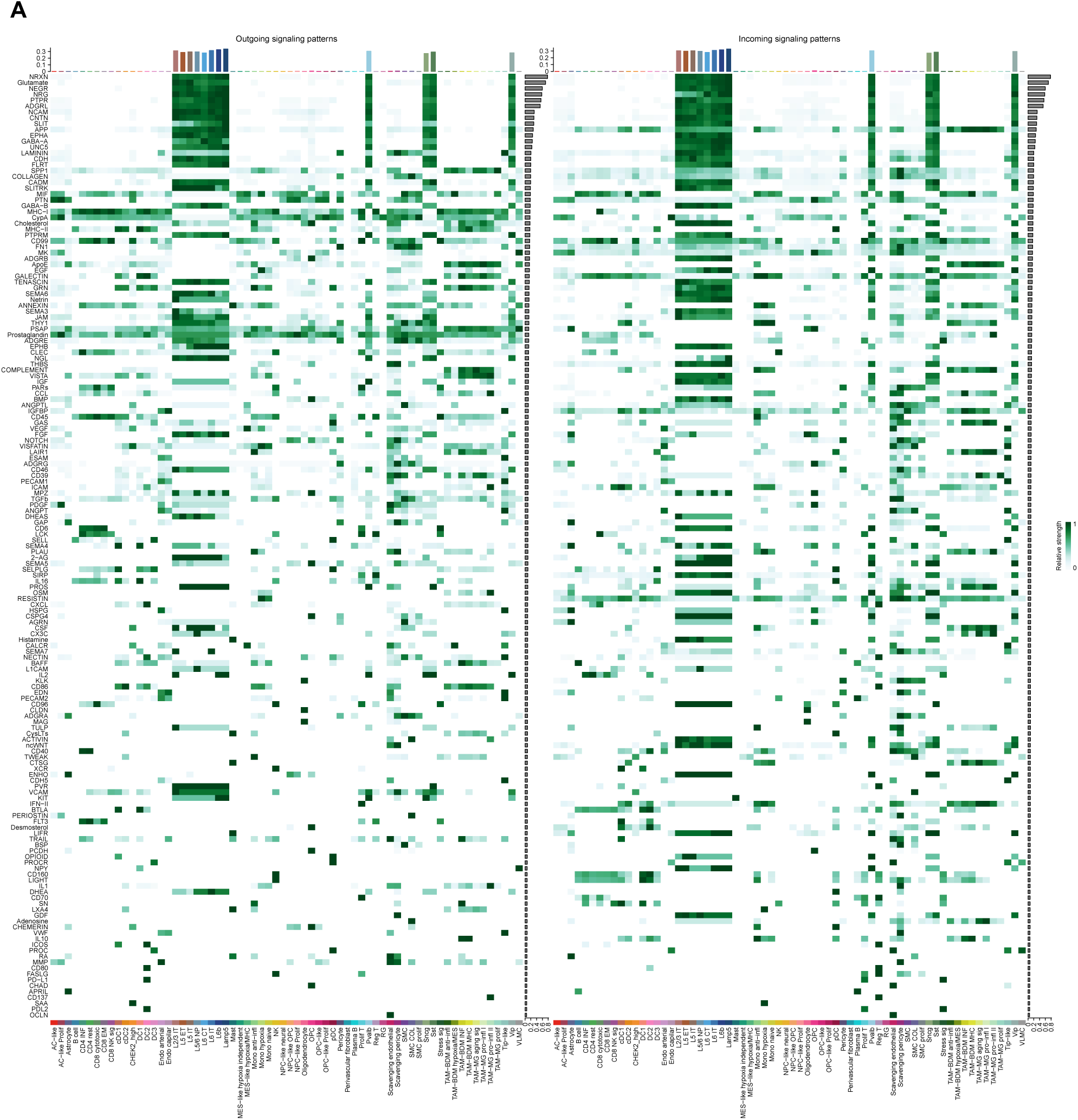
(A) Overview of all significant signaling networks. Outgoing signaling patterns are shown in the heatmap on the left and incoming signaling patterns are shown on the right. The relative strength of the signal is indicated in green.

## Supplementary Tables

**Table 1.**
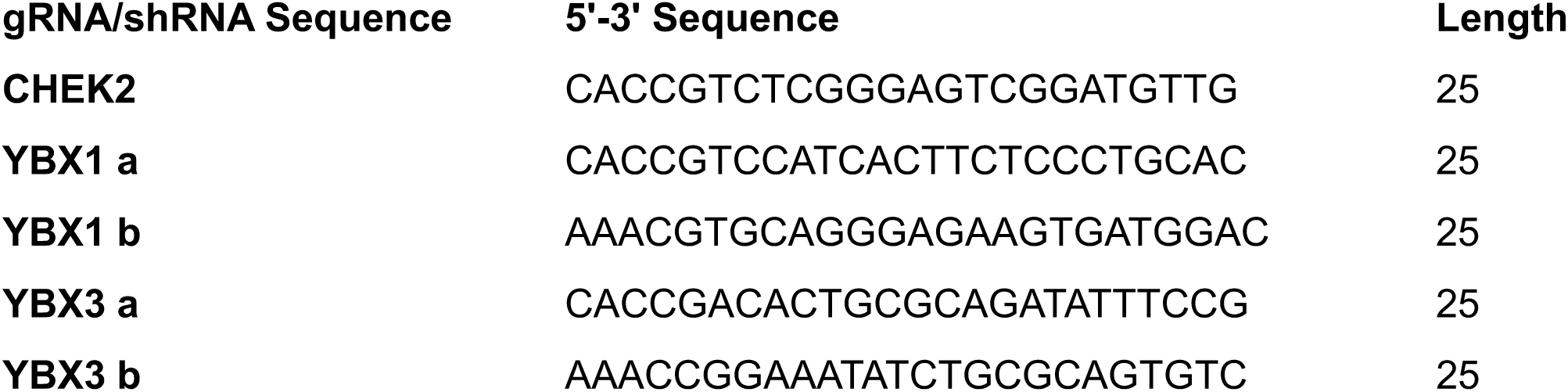
gRNA/shRNA sequences targeting human CHEK2, human and mouse YBX1, and human and mouse YBX3.List of guide RNA (gRNA) and short hairpin RNA (shRNA) sequences designed for targeting human CHEK2, human and mouse YBX1, and human and mouse YBX3. The sequences are presented in the 5’ to 3’ orientation, with a length of 25 nucleotides for each construct.

**Table 2.**
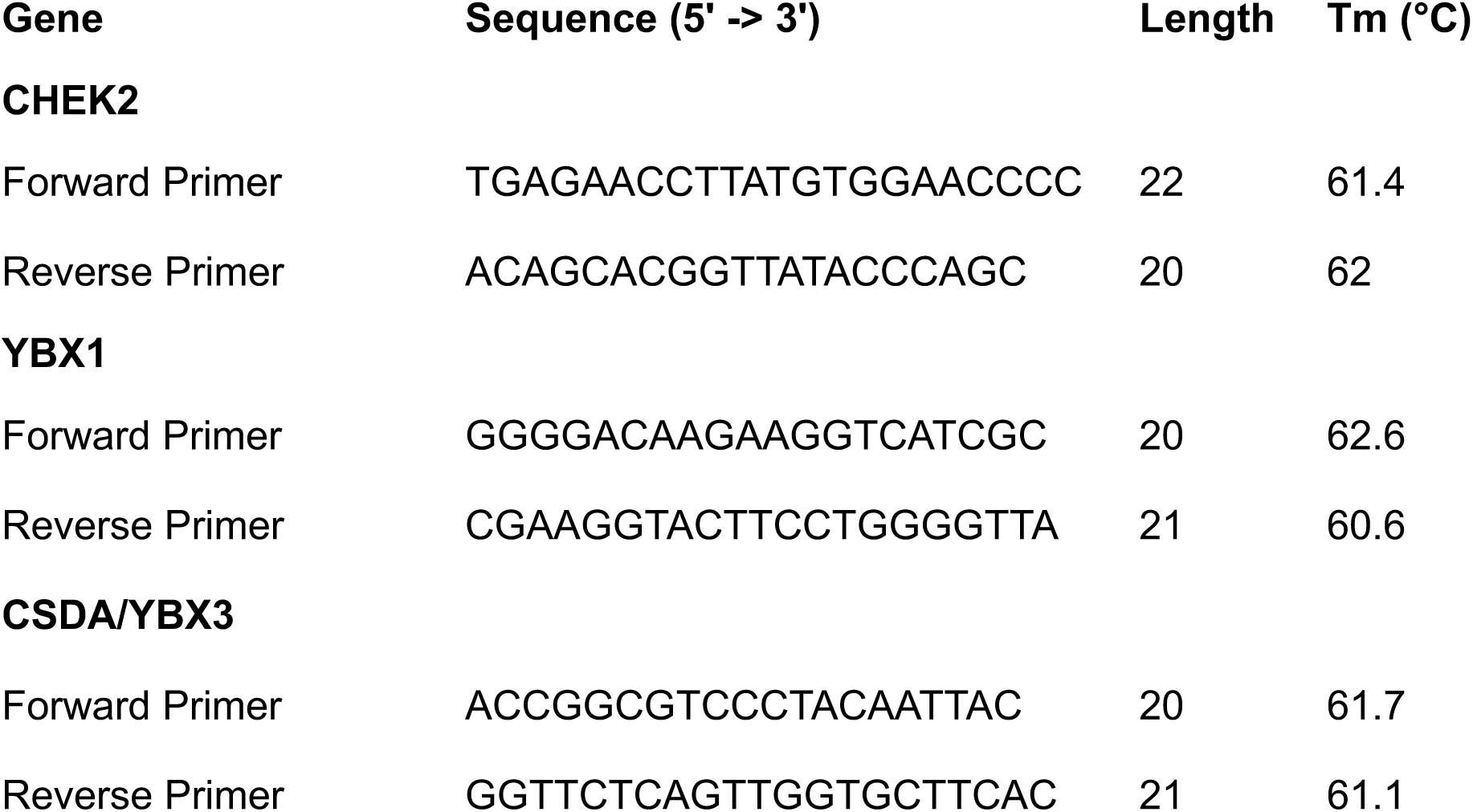
Primer sequences for CHEK2, YBX1, and CSDA/YBX3 in human and mouse. List of forward and reverse primers designed for PCR amplification of human and mouse CHEK2, YBX1, and CSDA/YBX3. Sequences are presented in the 5’ to 3’ orientation, along with their respective lengths (in nucleotides) and melting temperatures (Tm, in °C).

## Notes

### Competing Interest Statement

The authors have declared no competing interest.

